# Vivarium: an Interface and Engine for Integrative Multiscale Modeling in Computational Biology

**DOI:** 10.1101/2021.04.27.441657

**Authors:** Eran Agmon, Ryan K. Spangler, Christopher J. Skalnik, William Poole, Shayn M. Peirce, Jerry H. Morrison, Markus W. Covert

## Abstract

**Motivation:** This paper introduces Vivarium – software born of the idea that it should be as easy as possible for computational biologists to define any imaginable mechanistic model, combine it with existing models, and execute them together as an integrated multiscale model. Integrative multiscale modeling confronts the complexity of biology by combining heterogeneous datasets and diverse modeling strategies into unified representations. These integrated models are then run to simulate how the hypothesized mechanisms operate as a whole. But building such models has been a labor-intensive process that requires many contributors, and they are still primarily developed on a case-by-case basis with each project starting anew. New software tools that streamline the integrative modeling effort and facilitate collaboration are therefore essential for future computational biologists.

**Results:** Vivarium is a Pythonic software for building integrative multiscale models. It provides an interface that makes individual models into modules that can be wired together in large composite models, parallelized across multiple CPUs, and run with Vivarium’s simulation engine. Vivarium’s utility is demonstrated by building composite models that combine several modeling frameworks: agent based models, ordinary differential equations, stochastic reaction systems, constraint-based models, solid-body physics, and spatial diffusion. This demonstrates just the beginning of what is possible – future efforts can integrate many more types of models and at many more biological scales.

**Availability:** The models, simulation pipelines, and notebooks developed for this paper are available at the vivarium-notebooks repository: https://github.com/vivarium-collective/vivarium-notebooks. Vivarium-core is available at https://github.com/vivarium-collective/vivarium-core, and has been released on PyPI. The Vivarium Collective (https://vivarium-collective.github.io) is a repository of freely-available Vivarium processes and composites, including the processes used in Section 3. Supplementary materials provide with an extensive methodology section, with several code listings that demonstrate the basic interfaces.

## 1 Introduction

Our understanding of biological phenomena stands to be dramatically improved if we can adequately represent the underlying systems, mechanisms, and interactions that influence their behavior over time. Generating these representations, most commonly via mathematical and computational modeling, is made challenging by the complex nature of such systems. The most common modeling approaches today are statistical, extracting meaning from observational data by fitting functions such as regression, cluster analysis, and neural networks [1]. Although these models have been successful at approximating correlations among observed variables, the structures of the models are not easily interpretable, making it difficult to determine biological mechanism. In contrast, mechanistic models are designed to reproduce observed data by representing causality [2]. Thus, the mathematical form and parameters of mechanistic models are testable hypotheses about the system’s underlying interactions. In other words, mechanistic models can provide unique insights that can gather more evidence for or refute hypotheses, suggest new experiments, and identify refinements to the models. Some exciting progress has recently been made in combining both strategies [1,3,4].

Mechanistic models in computational biology have deepened our understanding of diverse domains of biological function, from the macromolecular structure and dynamics of a bacterial cytoplasm with atomistic models [5], the lysis/lysogeny switch of bacteriophage lambda with a stochastic models [6], bacterial growth in different conditions with constraint-based metabolic models [7], and cell-based models of quorum sensing in bacterial populations [8]. However, such models generally target a mechanism in isolation, with a single class of mathematical representation, and focus on a narrow range of resulting behavior. A logical next step in the development of computational biology is to combine these components and build upon their insights, so we can better understand how their mechanisms operate together as integrated wholes.

Integrative models combine diverse mechanistic representations with heterogeneous data to represent the complexity of biological systems. There have been several such efforts including the integrative modeling of whole-cells [9, 10], macro-molecular assemblies [11], microbial populations [12], and even some work towards whole-organisms [13]. They have shown some success in capturing the emergence of complex phenotypes – but many challenges remain to the extensibility of the resulting models and to their widespread adoption. This results in a loss of research momentum and an apparent ceiling on model complexity. The ideal model would be not only be integrative in terms of incorporating diverse mathematical approaches and biological functions, but also in terms of bringing together the vast scientific expertise across the globe. What is therefore required is a methodology that brings molecules and equations, as well as labs and scientists, together in this effort.

In this regard, software infrastructure can greatly facilitate the development of integrative models. Two major areas of development in this space are standard formats and modeling frameworks. Standard formats allow models to be shared between different software tools – just as HTML allows web pages to be viewed across multiple browsers and devices. Popular formats include FASTA for sequence encoding [14], Systems Biology Markup Language (SBML) for reaction network models [15], and Synthetic Biology Open Language (SBOL) for structural and functional information [16]. Model frameworks provide generic functions and objects that can be changed by users to write and simulate custom models within that framework. These include libRoadRunner for systems of differential equations [17], COPASI for stochastic simulations [18], Smoldyn for particle-based models [19], COBRA for constraint-based metabolic models [20], MCell for Monte Carlo [21], ECell for stochastic reaction-diffusion [22], cellPACK for spatial packing of molecular shapes [23], CompuCell3D for cellular Potts [24], PhysiCell for physical models of multicell systems [25], and BioNetGen for rule-based models [26]. However, committing to one approach can exclude insights that could be gained from others, and to date there is no established method to connect different approaches.

What we therefore need is a software solution for heterogeneous model integration, which allows many modeling efforts to be extended, combined, and simulated together. This hands control of model development to a community of users that can build their own modules and wire them together, rather than an in-house development team that maintains a rigid modeling environment. A shift to modular design can breed an ecosystem of models, which would interact with each other in large integrative models and possibly form symbioses of models that are known to work well together. The community of users would impose a type of selection pressure on these models to continually improve their accuracy and their reach.

This paper introduces Vivarium – software born of the idea that it should be as easy as possible for computational biologists to define any mechanistic model, combine it with existing models, and execute them together as an integrated multiscale model. It can apply to any type of dynamic model – ordinary differential equations (ODEs), stochastic processes, Boolean networks, spatial models, and more – and allows users to plug these models together in integrative, multiscale representations. Similar approaches have been developed for computer modeling of cyber-physical systems with Ptolemy II [27] and Modelica’s Functional Mock-up Interface [28]. Recently, related methods have also begun to be applied to synthetic and systems biology with modular environments such as Tellurium [29], CellModeller [30], and Simbiotics [31].

By explicitly separating the interface that connects models from the frameworks that implement them, Vivarium establishes a modular design methodology that supports flexible model development. Vivarium-core is a software library that provides the interface for individual models and for *composite models* – bundles of models that come wired together. Vivarium-core includes a discrete-event simulation engine, which takes input models, combines them, and runs them as coupled systems evolving over multiple time-scales. The plug-in system lowers the barrier to contribution, so that new users can more easily encode new processes and plug them into existing systems. We also present the Vivarium Collective, a registry of Vivarium-compatible models that can be imported into new projects, reconfigured, and recombined to generate entirely new models. The software has been designed to make it straightforward to publish Vivarium models as Python libraries on the Python Package Index (PyPI) to share with the community to plug into existing public or private models.

This paper is organized as follows: Section 2 provides a high-level overview of Vivarium’s features and introduces its terminology. A more detailed methodology is provided in supplementary materials, where we build an example system, starting with a deterministic model of unregulated gene expression, and then adding complexity through stochastic multi-time stepping, division, and hierarchical embedding in a shared environment. We note here that all of the examples we consider – both in the main text and supplementary materials – are available in Jupyter notebooks, designed so that readers can follow along in the code and execute the examples that are described in this paper. Using the tutorial and the corresponding Jupyter notebooks in combination, we hope that any prospective user of this software can get up-and-running very quickly. Section 3 demonstrates the integrative power of Vivarium, by combining several modeling paradigms into a composite model, including a genome-scale flux-balance model of metabolism, a stochastic chemical reaction network for gene expression and transport kinetics, and a solid-body physics engine for spatial multi-cell interactions. Finally, Section 4 discusses scalability and current limitations.

## 2 Vivarium overview

Vivarium does not include any specific modeling frameworks, but instead focuses on the interface between such frameworks, and provides a powerful multiscale simulation engine that combines and runs them. Users of Vivarium can therefore implement any type of model module they prefer – whether it is a custom module of a specific biophysical system, a configurable model with its own standard format, or a wrapper for an off-the-shelf library. The multiscale engine supports complex agents operating at multiple timescales, and facilitates parallelization across multiple CPUs. A glossary of key terms and definitions is included in supplementary materials; when first introduced here, they are shown in *italics*. All of the classes and methods are described in further detail in supplementary materials and are demonstrated in the supplementary Python notebooks.

Vivarium’s basic elements are *processes* and *stores* (Fig 1a,b), which can be thought of as software implementations of the update functions and state variables of dynamical systems. Consider the difference equation Δ*x* = *f*(*r, x*) · Δ*t*. A Vivarium store is a computational object that holds the system’s state variables *x*. A Vivarium process is a computational object that contains the update function *f*, which describes the inter-dependencies between the variables and how they map from one time (*t*) to the next (*t* + Δ*t*). Processes are configured by parameters *r*, which give the update functions a distinct shape of mapping from input values to output values.

**Figure 1:**
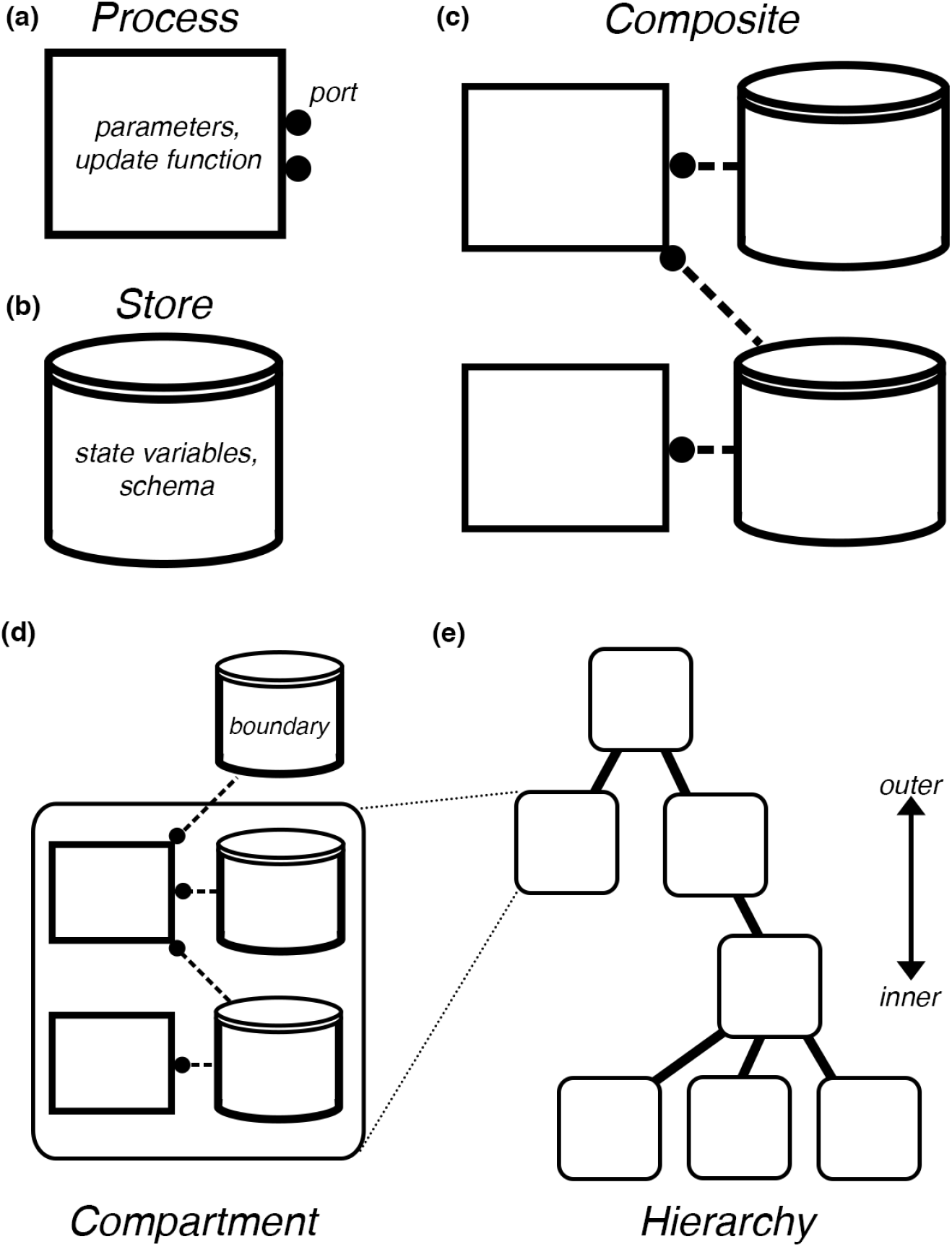
Vivarium’s model interface, illustrating the formal structure of the framework. (**a**) A *Process*, shown as a rectangular flowchart symbol, is a modular models that contain the parameters, an update function, and ports. (**b**) A *Store*, shown as the flowchart symbol for a database, holds the state variables and *schemas* that determines how to handle updates. (**c**) *Composites* are bundles of processes and stores wired together by a bipartite network called a *topology*, with processes connecting to stores through their ports. (**d**) *Compartments* are processes and stores connected across a single level. Processes can be wired across compartments through *boundary* stores. (**e**) Compartments are embedded in a *hierarchy* – depicted as a hierarchical network with discrete layers. Outer compartments are shown above and inner compartments below.

A *deriver* is a type of process for models that are not time-step-dependent (Fig 2d). For example, after the counts of molecules are updated by dynamic processes, it might be necessary to calculate the resulting change in total cell mass. A *mass deriver* would read all the updated molecular counts, and multiply by molecular weight to re-calculate cell mass. A second deriver could then read cell mass, and use it to update other values such as cell volume, length, and width based on a mathematical model of cell shape. Derivers run in a fixed order after the dynamic processes have finished. This allows users to implement a type of dependency graph – for example the mass deriver runs before the cell shape deriver. Derivers have been used to translate states between different modeling formats, implement lift or restriction operators to translate states between scales, and as auxiliary processes that offload complexity. In principle, derivers could also be used for statistical models that sample from an underlying distribution, or structural models that apply various constraints to determine spatial positioning of 3D shapes such as atoms or molecules into provided volumes [11,23].

**Figure 2:**
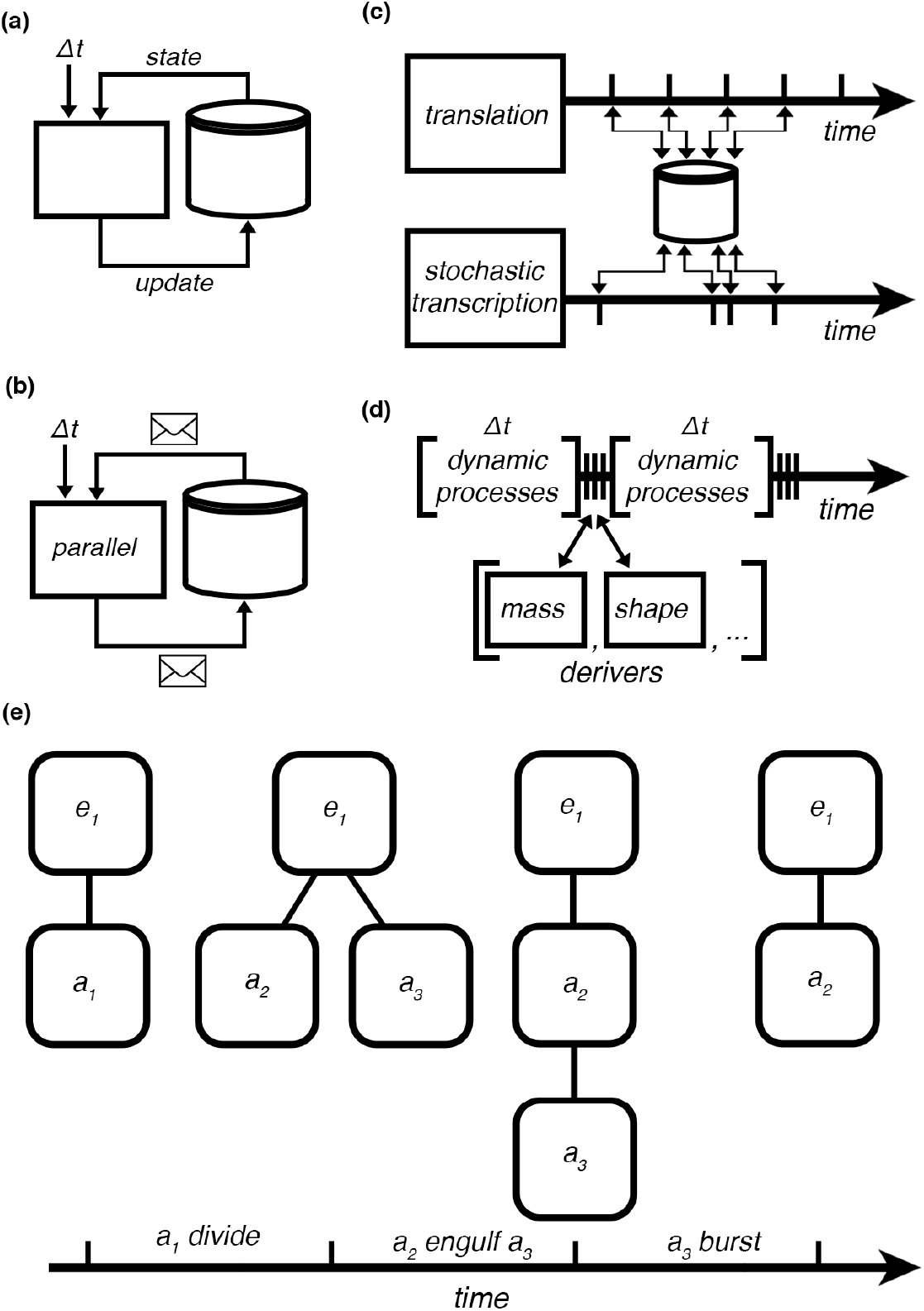
The vivarium engine takes the system and runs it forward in time. (**a**) A basic simulation loop has processes view the states through their ports, run their update function for a given time step, and return an update. (**b**) Any process can run in parallel on an OS process, which is placed on a long-running CPU thread, with state views and updates handled by message-passing. This allows for scalable simulation on computers with many CPUs. (**c**) Processes can run at different time scales, and using variable time steps. Here, a translation process is shown operating at fixed time steps, and stochastic transcription is operating at variable time steps determined based on the state of the system at each time step’s start. (**d**) Derivers are a subclass of Process, used for models that are not time-step-dependent. Derivers run in a fixed order at the end of every time step, and calculate additional states from states updated by the dynamic processes. In this figure a dynamic process runs for Δ*t*, followed by a mass deriver and a shape deriver in a fixed order. (**e**) A series of hierarchy updates depicts a compartment added by a *divide* update, then a compartment subsumed into a neighbor by an *engulf* update, then the engulfed compartment is deleted with a *burst* update. Other hierarchy updates include merge, add, or delete.

Processes include *ports*, which allow users to wire processes together through variables in shared stores. Variables in a store each have a *schema*, which declare the data type and methods by which updates to the variable are handled – this includes methods such as *updater*, for applying updates to the variables, and *divider* for generating daughter states from a mother state. A *topology* (short for “process-store interaction topology”) is a bipartite network that declares the connections between processes and stores, and which is compiled to make a *composite* model with multiple coupled processes. Ideal processes do not have hidden private states, and expose their states by externalizing them in stores. But sometimes private states are unavoidable, and could actually be used to improve performance since they do not have to synchronize. Externalizing state variables in stores allows other processes to wire to the same variables, which couples those processes – they read from and update these same variables as the simulation runs forward in time.

Processes can be connected across a hierarchical representation of nested compartments. Vivarium uses a bigraph formalism [32] – a network with embeddable nodes that can be placed within other nodes, and which can be dynamically restructured. This contrasts with the standard “flat” network that has all nodes at a single level, and with fixed connectivity. A *compartment* is a store node with internal nodes, which can include its own internal processes and the standard variable-containing stores (Fig 1d). A *hierarchy* is a place graph, or directory structure, which defines inner/outer nesting relations between compartments (Fig 1e). *Boundary* stores connect processes across compartments in a hierarchy – these make compartments themselves into pluggable models that can be embedded in a hierarchy. Just as with biological systems, compartments are the key to a model’s complexity – they organize systems into hierarchies of compartments within compartments, with modules that can be reconfigured and recombined.

In practical terms, this means that Vivarium facilitates collaborative model development by simplifying the incorporation of alternate sub-models. This allows users to 1) write their own processes, composites, and update methods, 2) import libraries with processes developed for different projects, 3) reconfigure and recombine existing processes, and 4) make incremental changes (add, remove, swap, reconfigure) and iterate on model designs that build upon previous work. Auxiliary processes are provided to offload complexity from the main processes.

The Vivarium *engine* is provided with the processes and a topology, it constructs the stores based on the processes’ declared schemas for each port, assembles the processes and stores into a hierarchy, and executes their interactions in time (Fig 2a). Vivarium aims to support large models with thousands of integrated mathematical equations. To accommodate these demands, Vivarium can distribute processes onto different OS processes (not to be confused with a Vivarium process) (Fig 2b). Communication between processes on separate OS processes is mediated by message passing with Python’s multiprocessing library. Simulations have run on Google Compute Engine node with hundreds of CPUs [33] – which scale the computation without any additional run time until there are as many processes as CPUs, after which some processes have to be run sequentially.

The engine is a discrete-event simulator – it advances the simulation forward by tracking the global time, triggering each process at the start of its respective time step, retrieving updates at the end of the time step, and passing these updates to the connected stores (Fig 2c). Processes can declare their own required time step, and can update their time steps during runtime to support adaptive time steps. The structure of a hierarchy is also dynamic and allows for stores, processes, and entire compartments to be created, destroyed, or moved during runtime. This allows for modeling important biological mechanisms that include forming, destroying, merging, division, engulfing, and expelling of subcompartments (Fig 2d).

## 3 Multi-paradigm composites

In this section, we demonstrate the power of Vivarium by applying it to complex, real-world examples. Specifically, we use Vivarium to integrate several modeling paradigms, building wrapper processes around existing libraries and wiring them together in a large composite simulation. COBRA is used for flux-balance analysis [20], Bioscrape is used to simulate chemical reaction networks [34], and pymunk is used as a solid-body physics engine for spatial multi-cell physics [35]. The implementation of each of these separate processes is briefly described here, and in greater detail in the supplementary materials; however, our primary aim in this section is to highlight the integrated, multi-paradigm model of an *E. coli* colony with many individual cells in a spatial environment, that collectively undergo a lactose shift in response to glucose depletion. For interested readers, we recommend the supplementary python notebooks, which show the incremental development steps, describe strategies for their integration, and display the resulting emergent behavior.

When grown in media containing the two sugars glucose and lactose, a colony of *E. coli* will first consume only the glucose until it is depleted; the colony will then enter a lag phase of reduced growth, which is followed by a second phase of growth from lactose uptake. During the glucose growth phase, the expression of the lac operon is inhibited while glucose transporters GalP and PTS are expressed. When external glucose is depleted, cells at first do not have the capacity to import lactose. The lac operon controls three genes: *lacY* (Lactose Permease) which allows lactose to enter the cell, *lacZ* (*β*-Galactosidase) which degrades the lactose, and *lacA* (Galactoside acetyltransferase) which enables downstream lactose metabolism. Once the operon is activated and proteins are expressed, the metabolism shifts to lactose and growth resumes. See [36] for a more comprehensive overview. For this example, a flux-balance model of *E. coli* is used to model overall cellular metabolism, while the details of the glucose-lactose regulatory, transport and metabolic circuit are represented by a chemical reaction network.

### 3.1 Individual paradigms

The individual processes were built with wrappers around existing modeling libraries, which were imported into this paper’s project repository and wired in a large integrative model. These are separately available for re-use in the libraries vivarium-cobra, vivarium-bioscrape, and vivarium-multibody. Supplementary material provides additional details on each of the processes, and shows each being run on its own.

#### Flux-balance analysis with COBRA

Flux balance analysis (FBA) is an optimization-based metabolic modeling approach that takes network reconstructions of biochemical systems, represented as a matrix of stoichiometric coefficients and a set of flux constraints, and applies linear programming to determine flux distributions [37]. FBA is made dynamic (called dFBA) by iteratively re-optimizing the objective with updated constraints at every time step [38]; these constraints change with environmental nutrient availability, gene regulation, or enzyme kinetics. We developed a Vivarium process that provides a wrapper around COBRApy [20], an API which can be initialized with genome-scale metabolic flux models [39]. The model used here is *iAF1260b*, which includes 2382 reactions and an objective that includes the production of 67 molecules [40].

#### Chemical reaction networks with Bioscrape

To add a chemical reaction network (CRN) model of transcription, translation, regulation, and the enzymatic activity of the lac operon and its resulting proteins, we turned to a published model [41], with many parameters from [42]. We converted this model to an SBML format using BioCRNpyler – an open source tool for specifying CRNs [43]. With the model in SBML, we ran simulations with a Vivarium process built with Bioscrape [34] – a Python package that supports deterministic and stochastic simulations.

#### Multicell physics with Lattice

The environment is implemented using a composite called Lattice from the vivarium-multibody library that includes *multibody* and *diffusion* processes. Multibody is a wrapper around the physics engine pymunk [35], which models individual agents as capsule-shaped rigid bodies that can move, grow, and collide. Multibody tracks boundary variables for each agent: location, length, width, angle, mass, thrust, and torque. It applies these to the physics engine, runs it, and returns a new location for each agent. Agents can update their volume, mass, and motile forces. Upon division, a custom divider is applied to the mother agent’s location of agents, so that daughters are placed end-to-end in the same orientation as the mother. Diffusion simulates bounded two-dimensional fields of molecular concentrations. Each lattice location holds the local concentrations of any number of molecules, and diffusion simulates how they homogenize across local sites. Agents can uptake and secrete molecules at their position in the field using the adaptor process “local field”.

#### Adaptor processes

Each process described above focuses on a different aspect of cellular physiology and behavior, applies a different mathematical representation, and formats its data by different standards. Integrating them requires data conversions and complex mapping of assumptions about their shared variables. This is achieved with a set of *adaptor* processes which convert between the expected units, reference frames, name spaces, data formats, and other representations. Adaptor processes required for integrating the Bioscrape and COBRA processes with the lattice composite include processes “local field”, “mass deriver”, “volume deriver”, and “flux adaptor” (described below).

### 3.2 Model integration and simulation results

The final composite model topology is shown in Fig 3a. The COBRA process’s metabolism and Bioscrape process’s gene expression and transport are coupled through a “flux bounds” store, with the Bioscrape process setting uptake rates with the flux adaptor process, and the COBRA process using them as flux constraints on the FBA problem. The Bioscrape process calculates deltas for each reaction’s substrates and products with its stochastic kinetic simulator. These deltas are then used by the flux adaptor to calculate a time-averaged flux, which it passes to the flux bounds store. The COBRA process uses these values to constrain the FBA problem’s flux bounds, which impacts the resulting calculated flux distribution and the overall growth rate. The Bioscrape and COBRA processes are also coupled through the transporter proteins, as well as internal metabolite pools that are built up by metabolism. Changes in the internal metabolite pools trigger the expression of genes (*lac* genes in this example), which in turn influence the kinetic transport rates. Thus, a causal loop is implemented between the Bioscrape and COBRA processes.

**Figure 3:**
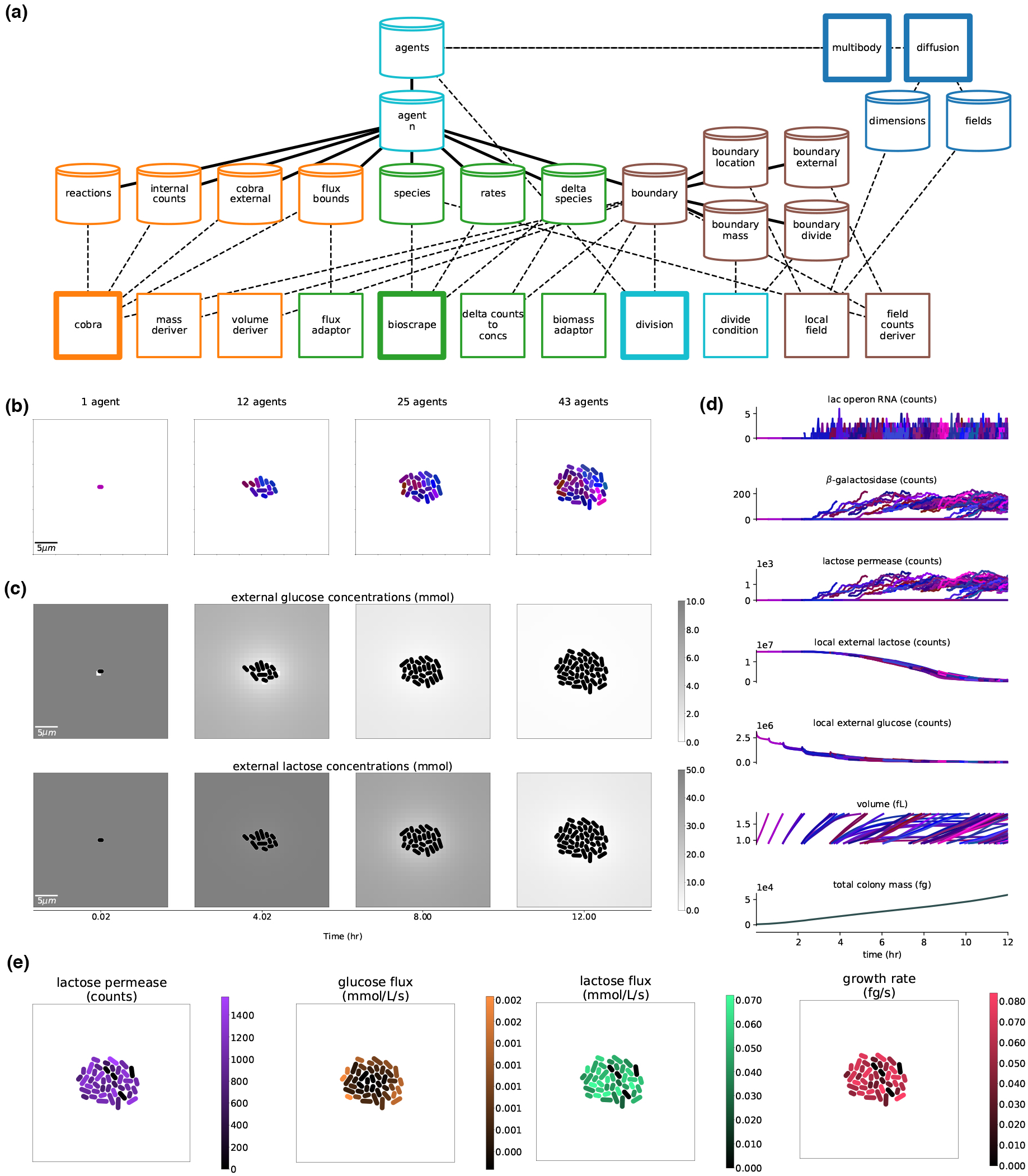
The full, integrated, stochastic version of the model in the Lattice environment. **(a)** Hierarchy and topology of the full composite model. The processes and stores are colored by paradigm, with orange nodes associated with COBRA, green nodes associated with Bioscrape, light blue with division, dark blue with multi-cell physics and chemical diffusion in the environment, and brown nodes associated with the boundary between cell and environment. Main processes are highlighted with a thicker outline. **(b)** Snapshots of the colony throughout a simulation. Cells colored according to phylogeny with similar colors indicating more closely related cells. **(c)** The external lactose and glucose concentration fields over the course of the simulation. **(d)** Multi-generational timeseries, with variables from all cells shown over time. These colors correspond to the phylogeny colors of (b). **(e)** Snapshots of the final simulation state (12.0*hr*), with various cell states tagged.

The stochastic versus deterministic versions of the composite require different adaptor processes for converting between counts and concentrations – for example, a “field counts deriver” process was required for the stochastic Bioscrape process to read external counts of glucose and lactose in a cell’s local environment based on the lattice resolution. The deterministic model runs much faster and for a larger local environment volume – as the local environment volume increases, this translates to many counts of glucose and lactose that the stochastic model has to simulate with the Gillespie algorithm, which is a time-intensive method. Thus, on its own, the stochastic model is limited to very small external environments and rapid depletion of nutrient. This challenge was overcome by partitioning the environment with the spatial processes described below. Future work could improve performance by passing reactions with small molecular counts to a stochastic simulator, and reactions with large counts to a deterministic simulator.

To model cell division, “division” and “divide condition” processes were added to read the agent’s individual cell mass, and trigger division when that agent reaches 2000*fg*. The mass deriver process takes the counts and molecular weights of these molecules (produced by the COBRA process), and calculates the total cell mass. The volume deriver process then calculates various cell shape properties of the cell from its mass, including volume, length, and width. These are used by the spatial environment. The divide condition process connects to the mass variable directly, and waits for it to cross a configured threshold value. When this threshold is passed, division performs the hierarchy update to terminate the mother agent and generate two daughters.

The environment consists of 2D arrays of concentration values for glucose and lactose, which set the local external environments for individual cell agents. The local field process converts COBRA process-generated molecular exchanges into concentration changes in spatial fields. The diffusion process takes the resulting fields, models diffusion between them, and updates the local external variables for each agent, so that agents only experience the concentrations at their given location. When agents reach the mass requirement for division they divide, after which their daughter cells are placed end-to-end, and their growth pushes upon neighbors with the multibody process. Thus, cell growth and division lead to the emergence of a colony with many individuals (Fig 3b).

The full model was used to simulate a glucose-lactose diauxic shift (Fig 3c-e). This simulation is configured with a low initial glucose concentration, so that the onset of lactose metabolism can be triggered within a few hours of simulation time. As the initial cell grows from a single agent through multiple generations (Fig 3b), it initially takes up glucose and not lactose (Fig 3c,d). The glucose at locations occupied by cells is locally depleted, but replenished by diffusion from neighboring locations (Fig 3c, top). The spatial variation in the environment leads to cells experiencing different local concentrations of glucose and lactose. In response, some lac operon RNA is expressed, due to stochasticity as well as in response to the local environment (Fig 3d). Expression of the lac genes allows lactose to be taken into the cells (Fig 3c, bottom), but with different levels of the lac proteins, leading to heterogeneous uptake rates for glucose and lactose as well as the growth rate (Fig 3e). By the end of the simulation, there is still slight glucose uptake; the colony is growing very slowly and is still shifting between growth phases, but lactose-driven growth has become the main driver of colony growth.

## 4 Discussion

This paper introduces Vivarium – a software tool for integrative multiscale modeling designed to simplify how we access, modify, build upon, and integrate existing models. We demonstrated the integration of several diverse frameworks including deterministic and stochastic models, constraint-based models, hierarchical embedding, division, solid-body physics, and spatial diffusion. Each process was developed and tested independently, and was then wired into a larger composite model that integrates these diverse mechanistic representations. While modular software exists that enables simulation of cellular biophysics and colony growth, such as CellModeller [30] or Simbiotics [31], or else allows for the assembly and integration of models based ordinary differential equations represented in SBML, such as Tellurium [29], Vivarium was designed with the requirements of whole-cell modeling in mind and as such is a far more general model integration tool. By enabling interfaces between a wide range of different models and supporting their interactions in a unified simulation, we believe that Vivarium will be broadly useful to diverse integrative models and a host of applications.

Users of any new model integration software should expect support for simulations of any size and complexity, as well as for arbitrary model simulators for different biological processes. Moreover, these simulations need to be accessible to as many scientists as possible. To meet these expectations, Vivarium was built to support three types of scalability: 1) scalable representation, 2) scalable computation, and 3) scalable development. Our discussion focuses on Vivarium’s strengths and limitations with regards to all three types of scalability, so that future iterations of model integration software can improve on the design.

### Scalable representation

A Vivarium model can be built with nested hierarchies that represent states and mechanism at multiple spatial and temporal scales. This multi-scale representation was demonstrated by running models at different levels of granularity – from stochastic CRNs operating on individual molecule counts, to genome-scale flux models of whole cell metabolic networks, to a rigid-body physics simulator with many individual cells. Hierarchies can be updated during run time to simulate behaviors such as division, merging, engulfing, and expelling – this enables the structure of the model to change, grow, and scale as new processes are launched and the total simulation state evolves. While agent-based, deterministic, stochastic and optimization-based models were demonstrated here, they represent only a single demonstration of the scalability of this framework. As another example, we recently applied Vivarium to simulate a multi-scale model o *E. coli* chemotaxis [44], and also used it to create the first “whole-colony” simulations of cellular growth on an agar plate [33]. In these previously-impossible, multi-cell simulations, every individual cell is an instance taken from the current version of the *E. coli* whole-cell model [10], and Vivarium integrates the behavior of these large-scale cell models as agents interacting across a lattice in a shared growth environment.

At present, there are no Vivarium processes to represent model uncertainty, which arises when some aspects of the system are not known such as the underlying mechanism or initial state, as can occur in stochastic models. In principle, if even a single variable in any component model has uncertainty associated with it, that uncertainty spreads to all variables in the other component models. Thus, it will be essential to develop specialized Vivarium processes to handle uncertainty in the future. One example might be a type of deriver, which would read the system state, evaluate uncertainty, and determine how to proceed – for example by triggering ensembles of processes to run, analyzing their resulting states, and using these states to determine the full system update.

### Scalable computation

Vivarium’s engine supports distributed simulation, with each model process running in parallel on its own OS process, and communicating with the engine through messages. The engine distributes the processes across a computer architecture with many CPUs, and runs them in parallel at their preferred timescales. One current limitation is that most simulators were not designed to be called iteratively in rapid succession with updated input states, and have to be re-initialized at the start of every time step; doing this can result in slower run time. One solution to this problem would be to optimize model simulators such that they can be called iteratively; such optimization already exists for libRoadRunner and COBRA.

A current limit on the size of parallel simulation stems from Vivarium’s use of the Python multiprocessing library. This allows it to run on a many-core computer, such as a Google Compute Engine (up to 224 cores at present). Parallel execution is made possible by handling communication between processes with message-passing so that each process can run on its own OS process. The same message-passing methodology can be used to distribute computation across many computers in a network, where tools such as mpi4py [45] would greatly extend Vivarium’s capabilities beyond a single compute engine.

Another current limitation is the requirement of Pythonic API, which renders non-Pythonic simulators harder to integrate. Requiring a Python API is not unreasonable, as Python has essentially become the standard language for scientific modeling, which means many simulators are already accessible. There are also libraries that support building Python APIs for other languages – for example, pybind11 simplifies how we can call C++ methods using Python. The next generation of model integration software might involve a lower-level language, and provide support for network-based APIs, which would allow processes to run in a distributed environment in their own native language.

### Scalable development

Vivarium was designed to support collaborative efforts, in which modular models are reused and recombined into increasingly complex computational experiments over many iterations with different contributors. Its emphasis on the interface between models simplifies the incorporation of alternate sub-models, which supports incremental, modular development. The Vivarium Collective is an early version of an online hub for vivarium-ready projects. When released on the Vivarium Collective, vivarium projects can be imported into other projects, re-configured, combined with other processes, and simulated in large experiments. We found that adaptor processes are very useful – they help convert the units, reference frames, name spaces, data formats, and other representations that are expected by the main mechanistic processes. Some limitations on development stem from Vivarium’s flexible design, which delegates most modeling decisions to its users. Good models can be combined with bad models, timescales can be mishandled, and there is no built-in framework to hand model uncertainty, as mentioned above. We believe that in a healthy ecosystem of models, a type of collective selection pressure that will drive quality up as users choose the best or most appropriate models for their needs.

### Conclusion

As multi-scale models such as these are further developed and expanded, we hope that Vivarium and its successors will be used to model cell populations, tissues, organs, or even entire organisms and their environments – all of which are based on a foundation of molecular and cellular interactions, represented using the most appropriate mathematics, and integrated together in unified composite systems. We look forward to seeing what the community will produce using these exciting new tools.

## Acknowledgements

We thank two anonymous reviewers for their thoughtful comments, which were significant in shaping the discussion section. We thank Jeremy Zucker for reporting a bug in the vivarium-cobra library, Richard Murray for discussion and feedback on the paper, and the Build-A-Cell community for organizing workshop discussions that helped motivate the use of Vivarium as a general model integration software tool.

## Funding

This work was supported by the Paul G. Allen Frontiers Group via an Allen Discovery Center at Stanford, as well as NIGMS of the National Institutes of Health under award number F32GM137464 to E.A., and NSF grant CBET-1903477 to W.P. The content is solely the responsibility of the authors and does not necessarily represent the official views of the National Institutes of Health.

## Notes

https://github.com/vivarium-collective/vivarium-core

https://github.com/vivarium-collective/vivarium-notebooks

### Competing Interest Statement

The authors have declared no competing interest.

### Summary of Updates

The manuscript's introduction and discussion have been re-written. Some of the examples have been pulled out to a supplement to reduce the manuscript's length. And there is added discussion of "deriver" processes, which are not time-dependent.

