## Supplementary Materials for "Vivarium: an Interface and Engine for Integrative Multiscale Modeling in Computational Biology"

.

**References** **26**

### 1 Introduction

This supplement to "Vivarium: an interface and engine for integrative multiscale modeling in computational biology" includes detailed examples, additional descriptions of libraries used in the main text, tables with key terms and Vivarium methods, and coding examples. Section 2 goes into detail as it builds an example system, starting with a deterministic model of unregulated gene expression, and then adding complexity through stochastic multi-time stepping, division, and hierarchical embedding in a shared environment. This example is built up incrementally, highlighting key features of the methodology that enable incremental construction of complex models. Section 3 introduces the individual libraries used to build the multi-paradigm composite in the main text. This includes the vivarium-cobra, vivarium-bioscrape, and vivarium-multibody libraries. The appendices include tools for debugging, performance profiling of different models, references for software availability with urls, tables with key terms and methods, and code examples for building Vivarium models.

#### 1.1 Vivarium's basic elements

Table S2: Elements of the Vivarium framework. Some of these elements have software analogs, which are referenced in monospaced `code` format.

| Term | Definition |
|---|---|
| Process | A modular sub-model that encodes a biological mechanism and can be composed with other processes to create a larger composite model. Particular process instances are created as subclasses of the `Process` class. |
| Store | A collection of state variables read by the processes, which contains methods for applying the processes' updates. State variables in stores that are shared by multiple processes are the only means of communication between those processes. A `Store` class instance is automatically constructed based on the processes' declared variables and their schema. |
| Port | A named connector on a process that gets connected to a store. Processes can declare one or more port, and the state variables they want to receive through these ports. |
| Schema | A state variable's declared data type, default value, and methods such as updaters and dividers, by which updates to the variable are handled. Schemas are declared by the processes' and are used to initialize stores at the start of a simulation. |
| Topology | Short for "process-store interaction topology", this is a bipartite network that declares how to connect processes to stores. It is declared as a Python dictionary for each process, with port names mapped to paths where stores are expected. |
| Composite | An integrated model with multiple processes whose connections to stores are specified by a topology. A `Composite` class has processes and a topology; it is passed to the engine to create the required stores. |
| Composer | A composer (subclass of `Composer`) generates composites by initializing a processes and specifying a topology for how they are wired together. |
| Deriver | `Deriver` is a subclass of `Process`. Deriver instances runs after the other processes and calculate additional state values from other available state variables – for example, concentrations from molecular counts. These are used to offload complexity from the dynamical processes. |
| Compartment | A store that contains inner stores and processes, rather than the standard store with state variables. Processes can connect to other compartments through boundary stores. |
| Hierarchy | A hierarchical network with nested stores, which can be thought of as a directory structure. A hierarchy can be updated during simulation runtime with update methods such as divide, move, and add. |
| Engine | vivarium-core's `Engine` class is a discrete-event simulation engine. It takes processes and a topology as input arguments, creates the stores based on the processes' ports schema, connects the processes, and runs the integrated model forward in time. |

### 2 Interface basics

This section introduces the elements and methodology of Vivarium by working through an example system of unregulated gene expression. We provide supplementary Jupyter notebooks (links are provided in supplementary materials 4.1) that implement each of the examples in executable code – technical readers are encouraged to have these notebooks open as they read through this section.

The guiding example separates gene expression into two processes: transcription – in which a gene is transcribed to form mRNA, and translation – in which the mRNA is translated to form a protein. We start by representing each of these functions as simply as possible with difference equations, and run them on their own. We then integrate them in a composite model and simulate them together. Next, we replace the deterministic transcription process with a stochastic process, and run a hybrid deterministic/stochastic simulation of gene expression that includes variable time steps. Finally, processes for cell growth and division are added, which allow the system to split into many separate agents that run in parallel in the simulation.

#### 2.1 Transcription process

The transcription process used here is called "Tx", and models mRNA synthesis from DNA. We define a system with a single mRNA species, $C$, transcribed from a single gene, $G$. The chemical reaction network (CRN) – which specifies reactants, products, and a set of reactions – takes the form:

$$G \xrightarrow{k_{tx}} G + C,$$

$$C \xrightarrow{k_{deg}} \emptyset.$$

This CRN can be simulated with the difference equation

$$\Delta C = (k_{tx}G - k_{deg}C)\Delta t,$$

with $C$ expressed from $G$ at rate $k_{tx}$, and degraded at rate $k_{deg}$. Quantities are in concentrations (mg/mL) – this comes in useful later when converting to concentrations from counts. For pedagogical reasons, this model ignores gene copy number, RNA polymerase abundance, strength of gene promoters, and availability of nucleotides – features that could potentially be added later to improve the model's realism.

##### 2.1.1 Ports.

An illustration of Tx is shown in Fig S2a. $G$ and $C$ are read through different ports, "DNA" and "mRNA", which are connected to two different stores, also called "DNA" and "mRNA". By default, a process's ports connect to stores that have the same name – the next subsection demonstrates more complex mappings. For a small model with only two variables splitting the variables into separate ports might seem excessive, but for larger models this is a useful design principle. Generally, port design should be used to organize variables by useful categories: locations such as cytoplasm, membrane, chromosome; molecule groups such as metabolites, proteins, chromosomes; or by other categories such as global variables, fluxes, concentrations.

Listing S1: Python implementation of the minimal transcription process, Tx. This demonstrates the Vivarium process interface. Defaults correspond to default parameter values which can be overwritten in the `Process` constructor. Unit conversions are supported by the pint library [7]. States are Python dictionaries which encode the file structure of a Vivarium model. Updates are similarly returned as in the same hierarchical dictionary format.

```
class Tx(Process):
    defaults = {
        'ktx': 1e−2,
        'kdeg': 1e−3}

    def ports_schema(self):
        return {
            'mRNA': {
                'C': {
                    '_default': 100 ∗ units.mg/units.mL,
                    '_updater': 'accumulate',
                    '_emit': True,
                }},
            'DNA': {
                'G': {
                    '_default': 10 ∗ units.mg/units.mL,
                }}}

    def next_update(self, time step, state):
        # Retrieve the state variables through the ports
        G = state['DNA']['G']
        C = state['mRNA']['C']

        # Run the model
        dC = (self.parameters['ktx']∗G − self.parameters['kdeg']∗C) ∗ time step

        # Return an update
        return {
            'mRNA': {
                'C': dC,
            }
        }
```

##### 2.1.2 Process interface.

Listing S1 shows Python code for the Tx process – an instance of the `Process` class. Making a process requires implementing the process interface, which involves the following constant and methods: 1) `defaults`: This class constant declares expected parameter names and values – even if only with empty values that get replaced upon initialization. The process constructor (`__init__`) accepts a list of parameters when a process is initialized, which override the defaults. 2) `ports_schema`: This method declares a process' ports ("RNA" and "DNA"), the variables that are accessed through those ports ($C$ and $G$), and their required schemas (Supplementary materials, Table S9). Any type of value can be used in the

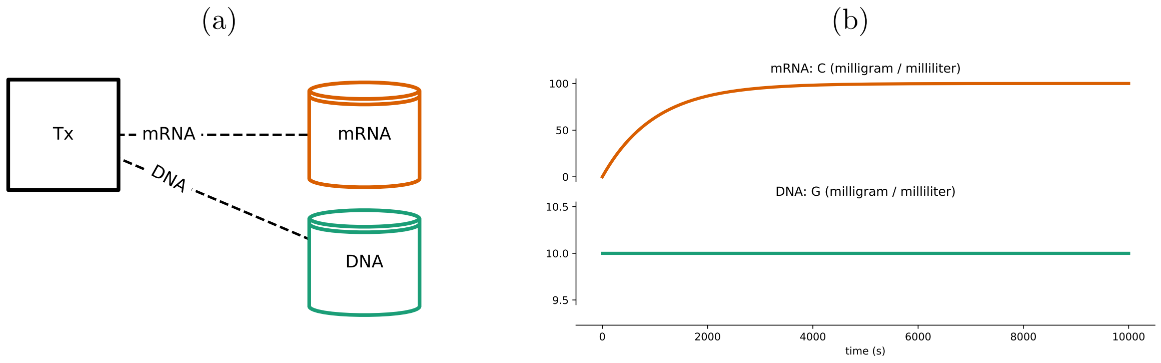

Figure S2: Tx – a transcription process. (**a**) The system's topology, with process Tx wired to two stores – DNA and mRNA – through ports of the same name. (**b**) Simulation output. The DNA $G$ remains fixed at its initial value, and the mRNA $C$ increases up to a steady state.

schema, such as integers, arrays, or more complex data structures. For novel data types, new schema methods such as updaters and dividers might be required – these are modular and can be defined by users. Updaters and dividers available with vivarium-core are listed in (Supplementary materials, Tables S10 and S11). 3) `next_update()`: This method contains the dynamical model. The steps of this method involves retrieving the variables through the ports, applying the encoded mechanism for the time step's duration, and returning the update for each port.

##### 2.1.3 Simulating a process.

The output of Tx is shown in Fig S2b. Individual processes can be run on their own by the simulation engine, which initializes the stores, runs the simulation, and saves the output. Each experiment is configured with an emitter, which logs the state of variables marked to emit during runtime – marking a variable to emit can be declared in the process' `port_schema`. If the emitter is connected a database (we use mongoDB), the saved data can be retrieved from the database for visualization and analysis at any time during or after a simulation run.

#### 2.2 Transcription/Translation composite

Next, we integrate the Tx transcription process with a translation process called "Tl", whose implementation is not shown here but is available in the supplementary notebook. Tl takes a similar form to Tx, but with protein $X$ translated from mRNA $C$ and degraded:

$$\Delta X = (k_{tl}C - k_{deg,X}X)\Delta t.$$

As before, units are in concentrations (mg/mL). As a simplifying assumption, translation rate considers ribosome availability, strength of ribosome binding to mRNA, availability of tRNAs and free amino acids to all be part of one lumped constant.

Listing S2: Python code for the TxTl Composer. This demonstrates the composer constants and interface methods. Processes are initialized in `generate_processes` and their ports are wired to stores by declaring a topology in `generate_topology`. Listing S6 in the Supplementary materials includes more advanced examples of `generate_topology`, including connecting ports to stores at different levels of the hierarchy, splitting a port into multiple stores, and connecting variables with different names.

```
class TxTl(Composer):
    defaults = {
        'Tx': {},
        'Tl': {}}

    def generate_processes(self, config):
        return {
            'Tx': Tx(config['Tx']),
            'Tl': Tl(config['Tl'])}

    def generate_topology(self, config):
        return {
            'Tx': {
                'DNA': 'DNA',
                'mRNA': 'mRNA'},
            'Tl': {
                'mRNA': 'mRNA',
                'Protein': 'Protein'}}
```

Fig S3a illustrates the composite model called “TxTl”, with Tx and Tl both wired to a shared store called mRNA. This couples the two processes, so that mRNAs synthesized by Tx impacts the expression of proteins by Tl.

##### 2.2.1 Composer interface.

`Composer` is a class that generates composites. A given composer’s inherited `generate()` method calls `generate_processes()` to construct the processes and `generate_topology()` to wire the processes to the stores. Then, it returns a composite that is ready for execution. Making a composer involves the following three class attributes: 1) Composers have their own `defaults` for parameters, which can override the default parameters for individual processes, thus providing easy access for parameter scans and learning algorithms to adjust the full composite’s behavior. 2) The `generate_processes()` method constructs a composite’s processes in a dictionary, which maps each of their names to the instantiated process objects. 3) The `generate_topology()` method returns a topology dictionary that declares how each process’s ports connect to stores in the hierarchy. The TxTl composite is specified in listing S2. Default parameters are empty so the processes will use their own defaults if none are supplied. Each process is constructed in `generate_processes()`, and wired together in `generate_topology()`. Tx’s DNA port maps to a store called “DNA”, Tl’s protein port maps to a store called “Protein”, and both Tx and Tl get wired to the same “mRNA” store containing the state variable “C”, thus coupling the two processes together.

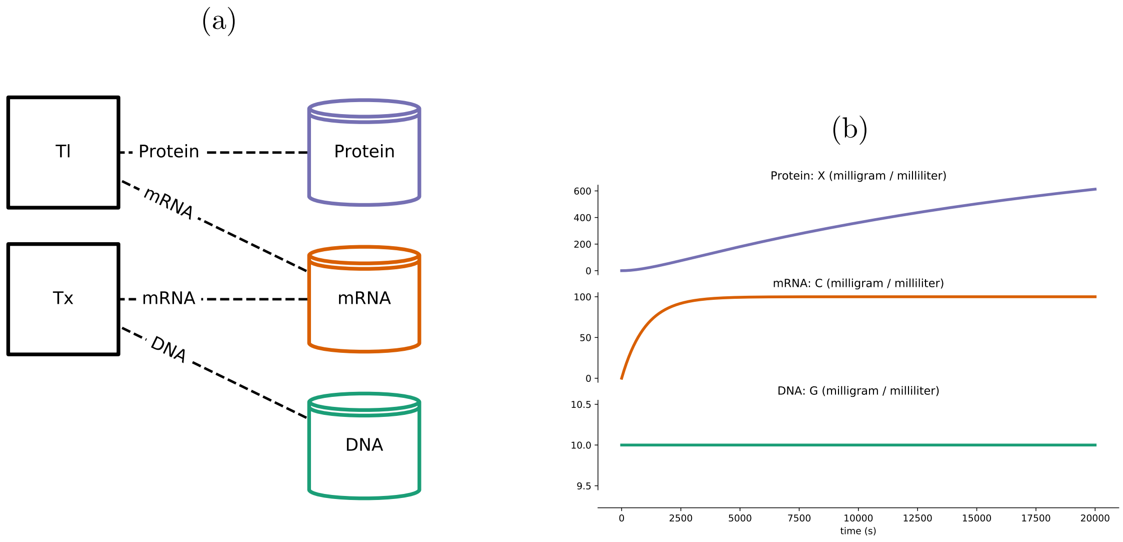

Figure S3: The TxTl composite with Tx transcription and Tl translation. (**a**) The model's topology and process declaration. Tx is the same as in Fig S2, and Tl is a new process with ports for mRNA and protein. Both processes are wired to the same mRNA store, thus coupling them together. (**b**) The simulation output of this model is the same as in Fig S2 showing DNA $G$ and mRNA $C$, but with added protein $X$ also being expressed.

##### 2.2.2 Using the vivarium engine

The vivarium engine runs TxTl to produce the simulation output shown in Fig S3b, where the mRNA reaches a steady-state as before while the protein concentration increases over time following a logistic-like curve.

Composite models are run with vivarium-core's `Engine` class. It is a discrete-event simulation engine that can accept either processes and a topology (the attributes of a `Composite` class) and generates the store hierarchy, or a pre-generated `Store` with the hierarchy already made. Some additional keyword arguments to engine let you pass in an initial state, emitter time steps, experiment name, and emitter type (for example print to console, save to RAM, save to database). Listing S3 shows how to take a composer, generate a composite, pass its processes and topology to the engine, and run a simulation.

#### 2.3 Adding complexity with a stochastic process

Importantly, Vivarium enables users to compare competing models of a given process, simply by exchanging one for the other and simulating the resulting behaviors. The interface introduced above makes it easier to define and integrate process modules – now we demonstrate how changes in the sub-models, building off prior model design, can provide additional model functionality.

We begin by replacing the deterministic transcription process, Tx, with a stochastic process called "stochastic Tx". The biological reasoning for this might be that in individual cells many genes are transcribed at low expression rates and synthesize small counts of mRNA, which leads to stochastic behavior. We use the Gillespie algorithm [4] – a discrete

Listing S3: Running a composite with `Engine`. This listing demonstrates how to initialize a composer by passing in parameters, how to generate a composite using a composer, and how to pass a composite's processes and topology into the simulation engine. The engine is then run for a length `total_time`, and its output data is retrieved from the emitter.

```
# define configuration, to override default parameters
config = {
    'Tx': {'ktsc': 1e−2},
    'Tl': {'ktrl': 1e−3}}

# initialize TxTl composer with the config
composer = TxTl(config)

# generate the composite, within initialized processes
composite = composer.generate()

# define an initial state, to override default values
initial_state = {
    'DNA': {'G': 10.0 ∗ units.mg / units.mL},
    'mRNA': {'C': 0.0 ∗ units.mg / units.mL},
    'Protein': {'X': 0.0 ∗ units.mg / units.mL}}

# initialize a simulation with the composite's processes and topology
experiment = Engine(
  processes=composite.processes,
  topology=composite.topology,
  initial_state=initial_state)

# run the experiment
total_time = 20000
experiment.update(total_time)

# get the simulation output
output = experiment.emitter.get_data()
```

and stochastic method for systems with few reactants – to simulate individual reactions. The Gillespie algorithm can be broken into two steps – one for calculating the time which elapses before an event occurs, and the other for determining the nature of that event. Stochastic simulations require variable time steps; for example, the distribution of time steps in a simulation using stochastic Tx is shown in Fig S4c.

##### 2.3.1 Derivers.

The Gillespie algorithm operates on molecular counts – every reaction event increases the counts of the products and decreases the counts of the substrates. Thus, the model needs to convert the molecular counts from the stochastic Tx process to concentrations for input to the Tl process. To perform this conversion, we add an auxiliary *deriver* process.

`Deriver` is a subclass of `Process`, but without a time components. These instances run after the dynamic processes, and in a fixed order like a dependency graph. For example, if

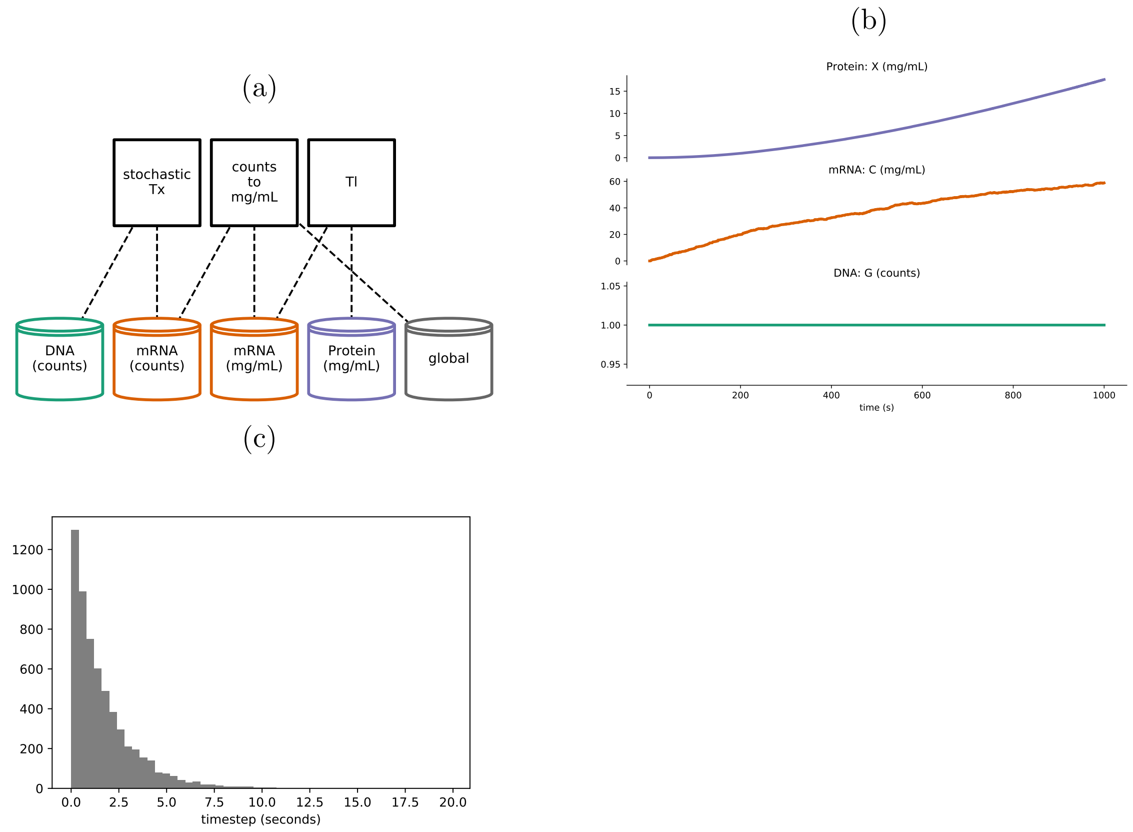

Figure S4: Stochastic transcription with deterministic translation. The stochastic process adjusts its time step based on the total propensity of the system, which is recalculated at every step. (**a**) The topology, showing the stochastic Tx process, the counts to mg/mL concentration deriver, and the deterministic Tl process. Counts to mg/mL is connected to a "global" store, which holds the volume variable required to calculate concentrations. (**b**) The simulation output shows stochastic dynamics of mRNA, and the impact of this stochasticity on the protein concentration. (**c**) Histogram shows the variable time steps resulting from running stochastic Tx on its own for $10,000$ seconds. Tl on its own runs at fixed 1 second intervals.

there were processes A and B with dt=1, C with dt=2, and two derivers D1 and D2, the run order would be something like:

$$t_0 : D1, D2$$

$$t_0 \rightarrow t_1 : A, B, D1, D2$$

$$t_1 \rightarrow t_2 : A, B, C, D1, D2$$

There are many uses for derivers. These could act as translators that adapt different modeling formats, lift and restriction operators that translate between scales, and helper processes to offload complexity.

The vivarium-core library provides several general-purpose derivers. In the current example we instantiate a deriver called "counts to mg/mL" in Fig S4 – this deriver calculates new concentrations from counts after every step.

##### 2.3.2 Multiple timescales.

For processes to operate at different timescales, the simulation engine handles updates on a per-process basis. At the start of each process' time step, the engine retrieves the process' required time step by calling its `calculate_timestep` method with the current state of the system. By default, the process returns a fixed time step that can be declared in their parameters, but stochastic Tx calculates a new time step by using the Gillespie algorithm. After retrieving the time step, the engine calls the process' `next_update` method with the current state of the system. When the system time reaches the end of the process' time step, it retrieves the update and applies it to the system state. This way, all processes can run at their preferred timescales. With the current version of the engine, the user needs to make sure the time steps of the processes are synchronized with each other to avoid numerical issues. Future versions can introduce a specialized adaptor process to handle the processes' time steps in a way that automatically ensures coordination.

#### 2.4 Hierarchical embedding

Up to this point, each model had a fixed number of processes and stores, with a fixed topology. In contrast, cell division requires a hierarchy with agents embedded in a shared environment, within which the cell agents can grow and divide. The hierarchy needs to launch new processes and stores for each agent created during runtime.

**Agents and environments.** When one compartment is nested in another, the inner compartment can be considered an agent and the outer compartment its environment. Coupling between an agent and an environment is supported by their processes sharing variables in boundary stores. For example, agent processes can update boundary variables required by environmental process such as agent volume, shape, motile forces, and uptake of molecules. Environmental processes can update the boundary conditions of agents' internal processes; for example, local molecular concentrations, and temperature. In the current example, a process called "colony volume" is added to the environment to calculate the volume of all the agents together (Fig S5a). This derived population-level state variable could in principle be used to drive other mechanisms – for example if simulated in a gut microenvironment, bacterial colony volume variable could be used to impact the host's digestion.

**Advanced process-store topology.** Up to this point, the system comprised a single compartment (i.e., a single cell). Accordingly, each topology specified a simple path from each port to a store in the same compartment – a "flat" network. Now, we model the environment as an outer compartment which can contain one or more cells. This requires advanced specifications in a composer's `generate_topology` that connect a processes' ports to stores further up or down the hierarchy. A port connects to a store in the hierarchy by specifying a path, which could go up the hierarchy to stores in outer compartments; or down the hierarchy to stores contained in inner compartments. `generate_topology` also supports splitting ports to draw from variables in separate stores, merging ports to draw from the same store, and aliasing names to variables to connect models with different variable names.

A few technical examples of these advanced topology methods are included in Supplementary materials section 4.6.

In the current example, the colony volume process reads the counts of all molecules through an “agents” store, which contains all of the individual agent instances – each with its own DNA, mRNA, and Protein, and global stores. Colony volume is configured to read the volumes in the individual global stores, and calculates total colony volume.

**Division.** To enable a compartment to divide during runtime, Vivarium provides a division process that is configured with a divide condition, which when true triggers division. A configurable condition means the process could be reused for more sophisticated cell models, for example based on the completion of chromosome segregation or the formation of a septum. Upon division, the mother’s variables’ states are divided between daughter agents based on those variables’ ’divider’ schema methods (Table S11).

For our example system, we initialize agents at 1000 fg, and trigger division when they double that mass. Fig S5c shows how the output over four generations of growth and division, starting off with a single stochastic TxTl instance that splits into two independent instances and then four and then eight – each of which exhibits its own distinctive behavior.

**Compartment hierarchy updates.** In biological systems, the nesting of compartments can be rearranged over time with behaviors such as engulfing, merging, and division. To support these behaviors, hierarchies in Vivarium can also be restructured during runtime. There are several built-in hierarchy update methods including move, add, delete, divide, and generate. Processes can trigger these in different combinations to generate a wide range of possible behaviors including merge, two neighboring compartments combine into one; burst, a compartment combines with its environment; engulf, one compartment is moved inside of a neighboring compartment; and expel, a compartment moves to be a neighbor of its outer compartment. All of these are available in vivarium-core.

Fig S5a shows the hierarchy after one division event, with two agents embedded in the top-level “agents” store. This requires the simulation to instantiate new agents – remove the mother agent, and generate two daughter agents with the mother’s composer, and an inherited state.

(a)

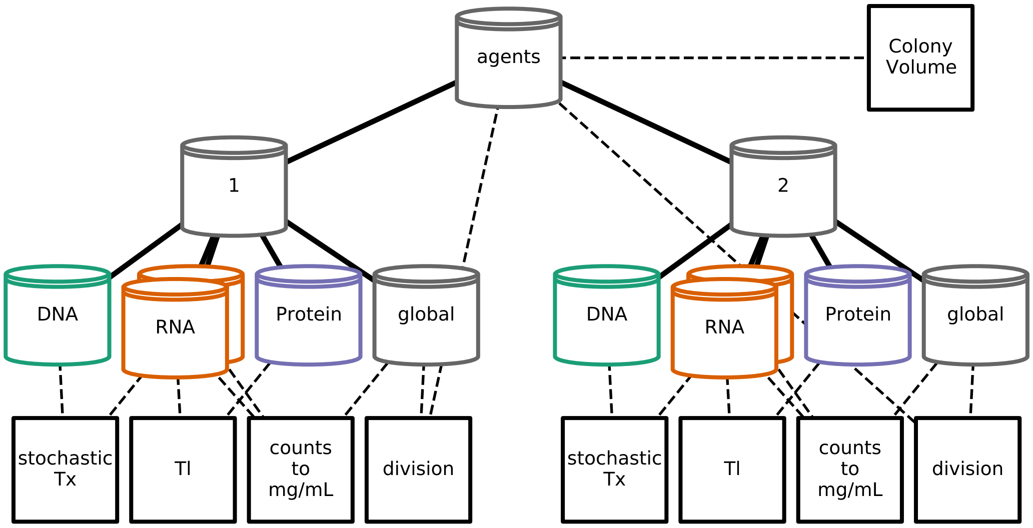

(b)

```
# make txtl composite with agent id "0"
agent_id = '0'
txtl_composer = TxTlDivision(agent_config)
agent0 = txtl_composer.generate(
    {'agents': agent_id})

# make a colony composite
colony_composer = ColonyVolume()
colony_composite = colony_composer.generate()

# perform merge
# agent0 in store "0", under "agents"
colony_composite.merge(
    composite=agent0,
    path=('agents', agent_id))
```

(c)

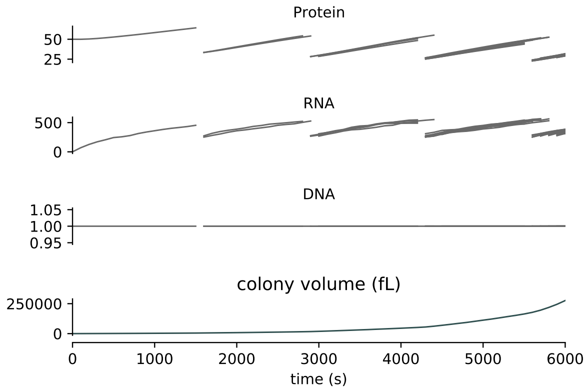

Figure S5: Hierarchical embedding and division. **(a)** Topology plot of the embedded hierarchy, with two cell agents. This plot was generated after one division event occurred, to illustrate how embedding works in Vivarium. Solid edges reflect hierarchy's directory structure, with "agents" at the top level. Dashed edges are topology connections between processes and the stores. **(b)** A script for making a composite model from a TxTl agent and a colony-level process to measure total colony volume. Agent 0 agent is placed within an 'agents' store, while Colony Volume is placed at the top level. The simplicity of the merge operation provides a flexible API for combining different models. **(c)** Simulation output of model over multiple generations of growth and division. Many instances run in parallel by the end of the simulation. The bottom plot shows the total colony volume, calculated from the full set of cell models within the environmental compartment.

### 3 Individual paradigms of Section 3

The main text demonstrates the integrative capabilities of of Vivarium by building a large multi-paradigm composite model of glucose-lactose diauxie in a colony of growing and dividing *E. coli* agents. The individual processes were built with wrappers around existing popular modeling libraries, which were imported into the glucose-lactose diauxie project and wired together into the large integrative model.

The individual processes are available for re-use in the Vivarium Collective: vivarium-cobra, vivarium-bioscrape, and vivarium-multibody. The PIPy versions used for this paper are vivarium-cobra==0.0.18, vivarium-bioscrape==0.0.0.7, vivarium-multibody==0.0.13. This section describes each of these libraries individually, shows their processes run on their own, and provides the parameters used to configure the example in Section 3.

#### 3.1 Flux-balance analysis with COBRA

Flux balance analysis (FBA) is an optimization-based metabolic modeling approach that takes network reconstructions of biochemical systems, represented as a matrix of stoichiometric coefficients and a set of flux constraints, and applies linear programming to determine flux distributions, for example those that maximize the production of biomass based on the known composition of metabolic end-products [6]. A strength of FBA is its capacity to simulate whole-network flux distributions using a minimal set of parameters. FBA is made dynamic (called dFBA) by iteratively re-optimizing the objective with updated constraints at every time step [12]; these constraints change with environmental nutrient availability, gene regulation, or enzyme kinetics. Many useful tools related to building and simulating FBA models have been developed and made freely available in the COBRA toolbox, which is also available in python as COBRApy [2].

For this work, we developed a Vivarium process that provides a wrapper around COBRApy, and is located in the vivarium-cobra library (Fig S6a). This process, called "COBRA", can be initialized with a BiGG model from the BiGG model database [5]. BiGG models are genome-scale metabolic models, which are available for dozens of *E. coli* strains, as well as many other cell types. The model used here is *iAF1260b*, which includes 2382 reactions, controlled by 1261 genes, and with an objective that includes the production of 67 molecules.

For purposes of integration with Vivarium, we pass the COBRApy results into internal metabolite pools that are available for other processes to utilize. The COBRA process includes a "flux bounds" port, which allows other processes to dynamically modify the flux constraints on the FBA problem. Accordingly, some additional processes were developed so that the COBRA process could support dFBA (shown in Fig S6a). These processes include "local field" to model the external environment with dynamic molecular concentrations, and "mass deriver" to convert the internal metabolite counts into a total mass.

Thus, when COBRA is run as a composite with the local field and mass deriver processes (Fig S6b), it takes up metabolites from the environment and grows its internal pools of metabolites, exponentially increasing in mass and reproducing the expected 40 minute doubling time in minimal glucose media.

(a)                (b)

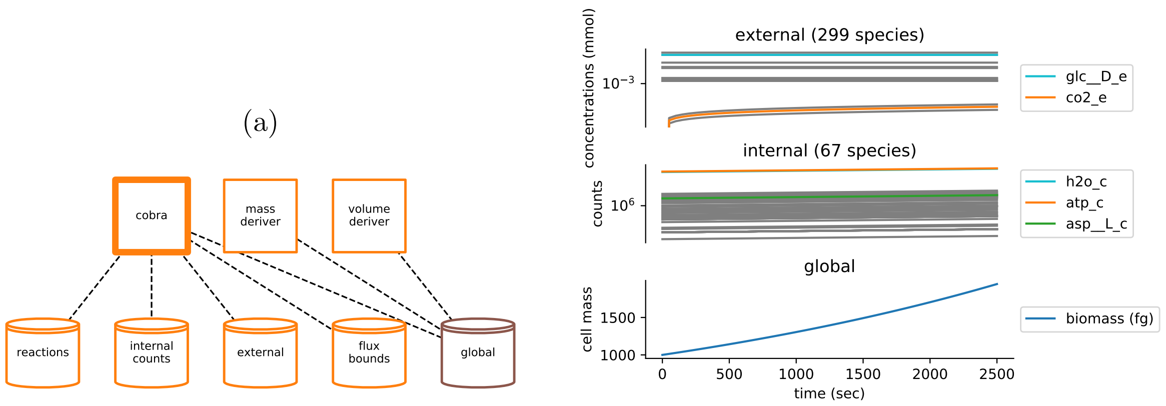

Figure S6: Demonstration of the COBRA composite's dynamic FBA. **(a)** Topology of the COBRA composite, which models metabolism with the COBRA process, and additional processes "local field" and "mass deriver" and help make the system into a dynamic FBA with an external environment, and total compartment mass based on individual metabolite pools and their molecular weights. The COBRA process has ports connected to "reactions", "internal counts", "external" (which are in concentrations), "flux bounds", and "global". **(b)** Simulation of a COBRA composite model, configured with BiGG model *iAF1260b*. Environmental concentrations (top) and internal molecule counts (middle) are plotted in log-scale due to the wide range across molecular species. The internal metabolites are multiplied by their molecular weight and summed to get total biomass (bottom).

| COBRA process | | |
|---|---|---|
| Parameter | Value | Description |
| BiGG model | $iAF1260b$ | metabolic model passed into the COBRA process, from [3] |
| time_step | $60s$ | fixed process time step |

Table S3: Processes parameters for vivarium-cobra version 0.0.18.

#### 3.2 Chemical reaction networks with Bioscrape

To add a CRN network model of transcription, translation, regulation, and the enzymatic activity of the lac operon and its resulting proteins, we turned to a published model [9], with many parameters from [13]. Our model includes all the same features, except for the addition of different combinatorial conformations of the lacR repressor binding to the lac operon. We converted this model to an SBML format using BioCRNpyler – an open source tool for specifying CRNs [8]. With the model in SBML, we were ran simulations with a Vivarium process built with Bioscrape (Bio-circuit Stochastic Single-cell Reaction Analysis and Parameter Estimation) [11] – a Python package that supports deterministic and stochastic simulations. The "Bioscrape" process is available at the vivarium-bioscrape library. Its topology can be seen in Fig S7a.

Running the lac operon CRN model in isolation shows expected behavior (Fig S7b), with glucose initially being taken up from the environment while lactose is not. Once external glucose is depleted, the lac genes are expressed, concentrations of $\beta$-Galactosidase and lactose permease rise, and lactose is brought into the cell and degraded. This is all done smoothly, with continuous dynamics (Fig S7b, left). Using the Bioscrape process also facilitates a stochastic simulation of this CRN (Fig S7b, right) the results of which show lac operon RNA expressed via a randomly-occurring transcription event, followed by expression of the lac proteins, and enabling subsequent import of lactose. For the stochastic model, external nutrients can only exist in a small external environmental volumes – large environments make for large nutrient counts, which slows the stochastic simulator drastically and makes simulations unfeasible. This limitation is corrected by the integrated model, which introduces many separate external locations for nutrients.

The Bioscrape process includes ports for "species", "delta_species", "rates", and "global". All the individual molecular species (including genes, RNA, and protein) connect to the the "species" port; "delta_species" is an output of the process that holds the changes (deltas) to the species in a given timestep, "rates" are instantaneous rates determined by the simulator, and "global" reads the "volume" variable.

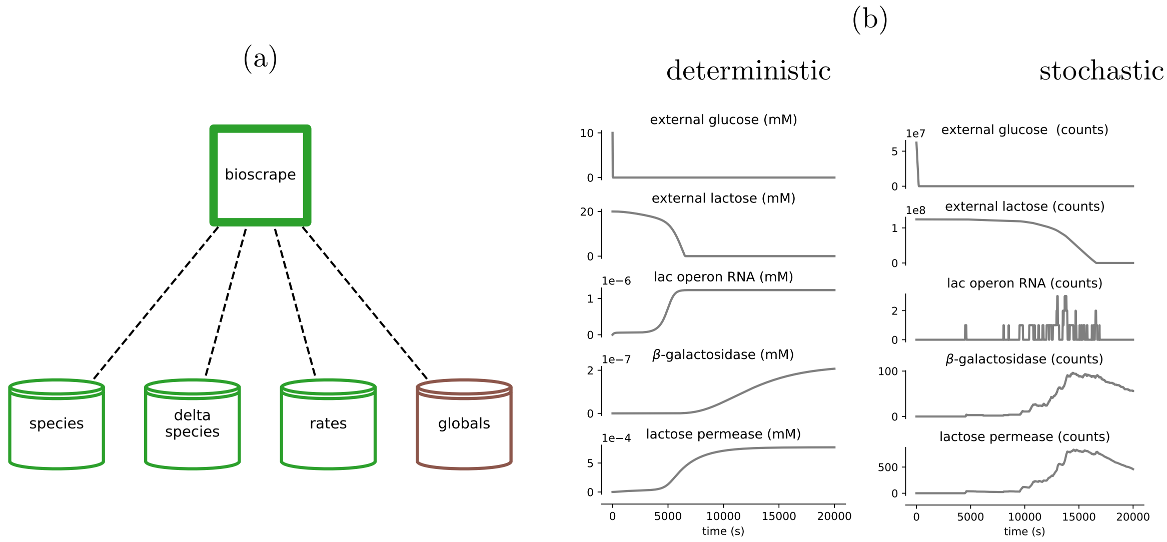

Figure S7: Demonstration of the Bioscrape process on its own. **(a)** Topology of the Bioscrape process, which models a CRN with deterministic (ODE) and stochastic (Gillespie) simulators. Bioscrape has ports connected to "species", "delta species", "rates", and "global". **(b)** Simulation of the Bioscrape process configured with a lac operon model. On the left – a deterministic simulation models the smooth dynamics of molecular concentrations. On the right – A stochastic simulation models the discrete events with molecular counts.

| Bioscrape process: deterministic model | | |
|---|---|---|
| Parameter | Value | Description |
| stochastic | $False$ | use Bioscrape's stochastic simulator |
| initial_volume | 1 | compartment volume for the simulator |
| internal_dt | 0.01 | internal time step with which to run the simulator |
| time_step | $10s$ | fixed process time step |

| Bioscrape process: stochastic model | | |
|---|---|---|
| Parameter | Value | Description |
| stochastic | $True$ | use Bioscrape's stochastic simulator |
| safe_mode | $True$ | use Bioscrapes safe model interface |
| initial_volume | 1 | compartment volume for the simulator |
| internal_dt | 0.01 | internal time step with which to run the simulator |
| time_step | $10s$ | fixed process time step |

Table S4: Processes parameters for vivarium-bioscrape version 0.0.0.7.

#### 3.3 Multicell physics with pymunk and field diffusion

With individual cells being represented by the Bioscrape and COBRA processes, our final step was to model a spatial environment in which these cells can grow, divide, and interact – through physical forces as well as by uptake and secretion of molecules in a shared chemical milieu. The environment is implemented using a composite from the vivarium-multibody library called "lattice", which consists of two processes: "multibody" and "diffusion" (Fig S8a).

The multibody process is a wrapper around the physics engine pymunk [1], which can model individual cell agents as capsule-shaped rigid bodies that can move, grow, and collide. Multibody tracks the following boundary variables for each agent: location, length, width, angle, mass, thrust, and torque. The physics engine applies these variables for the update time step, and returns a new location for each agent. Agents can update volume, mass, and motile forces, thus impacting their movement in the environment. Upon division, the `daughter_location` divider is applied to the location of agents, so that when they divide the daughters are placed end-to-end in the same orientation as the mother.

The diffusion process simulates bounded two-dimensional fields of molecular concentrations. Each lattice site $(x, y)$ holds the local concentrations of any number of molecules, and diffusion simulates how they homogenize across local sites. Each agent can uptake and secrete molecules at its position in the field. The implementation uses an adaptor process called "local field", which converts exchanged molecules from the given agent to concentrations at the agent's location.

Fig S8b shows the lattice composite simulated with minimal grow/divide agents. A single initial agent grows and divides to form colonies of many minimal agents in the environment – as they grow, they push against each other via the multibody process, and the colony increases in volume. The agents shown in this minimal simulation do not take up molecules. Therefore, in order to demonstrate the diffusion process, we initialized the system with a concentration gradient, which lessens over time.

| Lac operon expression and transport kinetics | | |
|---|---|---|
| Parameter | Value | Description |
| Translation and complexation of $\beta$-Galactosidase and lacP | | |
| k_tl_beta_Gal | 9.4/60 | translation rate of $\beta$-Galactosidase |
| k_tl_lacP | 18.8/60 | translation rate of lacP |
| BGal_ tetramerization | 1000 | complexation rate of $\beta$-Galactosidase, assumed to be fast and irreversible |
| Degradation of lactose with proportional Hill function | | |
| BGal_vmax | 300 | rate constant of lactose degradation to allolactose, catalyzed by $\beta$-Galactosidase |
| Bgal_Kd | 1.4 | dissociation constant |
| Import of glucose with proportional Hill function | | |
| GluPermease_vmax | 301 | rate constant |
| GluPermease_Kd | 0.015 | dissociation constant |
| LacPermease_ reverse_vmax | 71.38/60 | rate constant for reverse reaction |
| LacPermease_Kd | 14.62 | dissociation constant |
| Protein degradation with mass action | | |
| kdeg_mRNA | 0.47/60 | mRNA degradation rate |
| kdeg_prot | 0.01/60 | protein degradation rate |
| k_dilution | 0.02/60 | dilution rate for deterministic model, corresponds to a doubling time of 30 minutes. |
| stochastic model parameters | | |
| Bgal_Kd | 84310 | dissociation constant for stochastic model |
| GluPermease_Kd | 9033 | dissociation constant for stochastic model |
| LacPermease_Kd | 8800000 | dissociation constant for stochastic model |

Table S5: Model parameters for deterministic and stochastic models of the lac operon. These are a mixture of mass action and Hill function propensities. They were used with BioCRNpyler [8] to generate the SBML model and save '.xml' files, which are read by the Bioscrape Vivarium process. The files, LacOperon_deterministic.xml and LacOperon_stochastic.xml are available in the vivarium-notebooks repository.

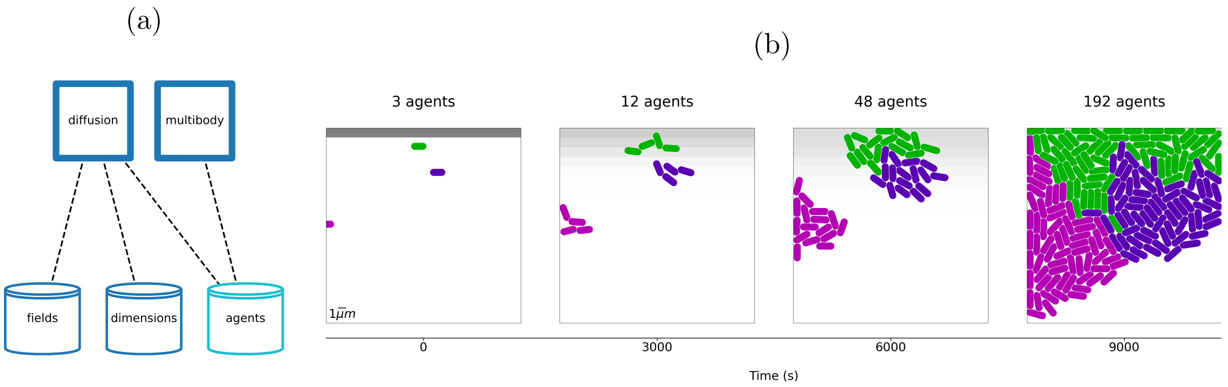

Figure S8: Demonstration of Lattice composite. **(a)** Topology of lattice composite, with a diffusion process and a multibody process, and ports connected to "fields", "dimensions", and "agents" stores. **(b)** Three grow/divide agents are initialized in the lattice. As the agents grows and divide, the multibody process simulates volume exclusion, which pushes their neighbors away and grows the colony. In this particular case, the agents do not exchange molecules with the external field, but diffusion can be seen by the spread of molecular concentrations initialized at the top row of the field.

| Diffusion process | | |
|---|---|---|
| Parameter | Value | Description |
| bounds | $(30\mu m, 30\mu m)$ | (x, y) dimensions of the environment. |
| n_bins | $(30, 30)$ | Number of discrete bins in the (x, y) dimensions. |
| depth | $0.5\mu m$ | The depth of the environment. Shallow environment deplete faster. |
| concentrations | $glucose : 10mmol$ <br> $lactose : 50mmol$ | Initial concentrations of the environmental fields. |
| diffusion | $0.02\mu m^2/s$ | Diffusion rate constant for all molecules. |

| Multibody process | | |
|---|---|---|
| Parameter | Value | Description |
| bounds | $(30\mu m, 30\mu m)$ | (x, y) dimensions of the environment. |
| jitter_force | $1 * 10^{-4} pN$ | A random force applied to each agent to simulate Brownian motion. |
| agent_shape | *segment* | The shape of the agents. Segment shapes are like rectangles with rounded ends – a 2D capsule. |

Table S6: Processes parameters for vivarium-multibody version 0.0.13.

### 4 Implementation details and support

#### 4.1 Software availability

The Vivarium Collective is a hub for Vivarium modules, and includes all of the libraries used for this project – https://vivarium-collective.github.io. These libraries are listed in Table S7. All of the notebooks are available for Jupyter on the vivarium-notebooks repository. For easier accessibility, the work described in this manuscript is also available on Google Colab (Table S8).

Documentation for vivarium-core is available at https://vivarium-core.readthedocs.io/en/latest/. The Vivarium Collective is a hub for Vivarium projects https://vivarium-collective.github.io.

Table S7: Vivarium libraries used in this project. These are all freely available for open development at the Vivarium Collective, and as pip-installable libraries on PyPI.

| Library | Description |
|---|---|
| vivarium-core | The interface classes and multiscale simulation engine. |
| vivarium-notebooks | The repository developed for this paper, with Jupyter notebooks and Python files. |
| vivarium-bioscrape | For chemical reaction networks with SBML. |
| vivarium-cobra | For constraint-based models of metabolism, with BiGG model interface. |
| vivarium-multibody | Lattice composite model used for spatial multi-cell interactions. |

Table S8: Python notebooks demonstrating paper simulations. Jupyter notebooks are available on GitHub, and require local installation. Colab notebooks run online, and no local installation is required.

| Description | Link |
|---|---|
| Vivarium interface basics Jupyter notebook. | https://github.com/vivarium-collective/vivarium-notebooks/blob/main/notebooks/Vivarium_interface_basics.ipynb |
| Multi-paradigm composites Jupyter notebook. | https://github.com/vivarium-collective/vivarium-notebooks/blob/main/notebooks/Multi-Paradigm-Composites.ipynb |
| Building the lac operon models with BioCRNpyler Jupyter notebook. | https://github.com/vivarium-collective/vivarium-notebooks/blob/main/notebooks/Lac_Operon_CRN.ipynb |
| Vivarium interface basics Google Colab notebook. | https://colab.research.google.com/drive/1-j2OXsflV8PZMbi9xldrCLkZ9Fj-y9sm?usp=sharing |
| Multi-paradigm composites Colab notebook. | https://colab.research.google.com/drive/1aiJ6uNjeATP0JTedxoZuNrLf3IPnHLU8?usp=sharing |

#### 4.2 Tools for debugging

It is important to provide tools for debugging, so that users can more easily build models, find sources of error in the model, and feel confident in their results. Vivarium is a powerful tool to connect models together, but if the connection of processes' ports to stores is not done properly, the whole model would not work or generate spurious results. Therefore it provides some tools to aid in debugging the wiring of models.

There are two main tools at present. When initialized with processes and their topology, `Engine` checks whether the topology matches the ports structure. If all the ports are successfully maps, it generates the stores based on port schema specifications; if the ports are not all mapped, it quits and prints a message about which process and which of its ports were not successfully mapped by the user's topology declaration. This feedback can be used to ensure all ports are mapped to stores.

Another useful tool is a function called `plot_topology`, which is provided with vivarium-core (Fig S9). This function plots a composite's connectivity, with processes, stores and their connections. By seeing the connections visually, users can better track their model's structure and process interactions.

(a)

```
# import the plotting function
from vivarium.plots.topology import plot_topology

# initialize a composite
txtl_composite = TxTl()

# plot topology topology
fig = plot_topology(txtl_composite, settings={})
```

(b)

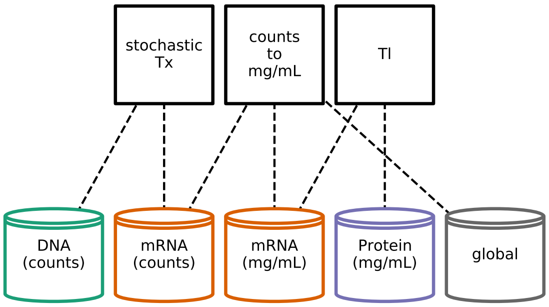

Figure S9: `plot_topology` is a useful tool for debugging by visualizing how processes and stores are connected. **(a)** Python script for generating a composite and plotting its topology **(b)** Example output, showing how three processes are connected to five stores.

#### 4.3 Performance

**Vivarium overhead.** To evaluate the overhead that Vivarium adds to runtime, we profiled simulations with models of varied complexity (can be found at `https://github.com/vivarium-collective/vivarium-core/blob/master/vivarium/experiments/profile_runtime.py`). These simulations can be systematically varied by the number of processes, the number of stores, the number of ports per process, the number of variables per port, and the hierarchy depth. Each run is evaluated by the cumulative *simulators time* and cumulative *Vivarium backend time*. This allowed us to measure the Vivarium backend costs scale relative to time in simulators as composite complexity increases.

When running a simulation without parallel processes, the simulator runtime scales linearly with the number of processes – this is the cost of the actual relevant scientific computation. The Vivarium backend also scales linearly with added processes, but at a rate smaller than an expected simulator' runtime – this cost comes from the engine's calls to each processes, and the application of each process' updates to the stores. The Vivarium backend also scales linearly with the number of ports for each process, with the number of variables that go through each port, and with the depth of the store hierarchy. We take this to mean that Vivarium users can engineer faster simulations by reducing the number of ports, reducing the number of variables, and reducing the hierarchy depth – there is room for optimizing some of these overhead costs, to reduce linear time $O(n)$ and possibly to constant time $O(1)$. But given this user discretion on composite wiring, we believe that performance limits on simulation will be driven primarily by the scientifically-meaningful simulation code, not any overhead from Vivarium.

**Parallel processes.** In a separate paper, we used Vivarium to run large simulations on a Google Compute Engine virtual machine (VM) with 96 CPUs and 442 GB memory [10]. In the time since we ran those simulations, Google Cloud has started offering VMs with 224 CPUs, which would enable even larger simulations. To fully utilize these many-core machines, individual processes are run in parallel using Python's multiprocessing library. This allows the simulation to add process without any additional overhead for individual simulators, as they all run in parallel until the number of vivarium processes equals the amount of available CPUs – after this the runtime again increases linearly.

**Technical specifications.** The simulations shown in Section 3, and in supplemental Section 3 were run by executing the simulation functions in `bioscrape_cobra/simulate.py`, which can be found in the Vivarium-notebooks repository on GitHub. The simulations shown were run on a MacBook Pro, with a 3.1 GHz Quad-Core Intel Core i7 processor. The multi-paradigm composite simulation shown in Section 3 was completed in three hours.

#### 4.4 Emitters

There are different `Emitter` classes to save simulation output, which can be specified with arguments to `Engine`. `RAMEmitter` saves the simulation output to RAM – this is useful for small simulations that you might not need to return to. `DatabaseEmitter` saves simulation output to MongoDB. Other emitters can be developed by subclassing the `Emitter` class.

#### 4.5 Ports schema

Table S9 covers the different types of ports schemas, which are declared by the processes. Updater and divider schema have their own tables – S10 and S11 – with methods provided by vivarium-core. In addition to these provided methods, custom updaters and dividers can easily be defined by the user for any type of desired state.

Table S9: Schema. A process' `ports_schema` method declares the schemas for variables connected to each port. These are used to construct the stores, which apply the declared methods during runtime.

| Attribute | Method |
|---|---|
| _default | The default value of the state variable if no initial value is provided. This also sets the data type of the variable, including units. |
| _updater | How to apply state variable updates. Available updaters are listed in Table S10. |
| _divider | How to divide the state variable's values between daughter cells. Available dividers are listed in Table S11. |
| _emit | A Boolean value that sets whether to log this variable to the simulation database for later analysis. |
| _properties | User-defined properties such as molecular weight. These can be used for calculating variables such as total system mass. |

Table S10: Updaters available in vivarium-core. Updaters are methods by which an update from a process is applied to a variable's value. New updaters can be easily defined and passed into a port schema (see Listing S4).

| Name | Function |
|---|---|
| accumulate | The default updater. Add the update value to the current value. |
| set | The update value becomes the new current value. |
| merge | Update an existing dictionary with new values, and add any newly declared keys. |
| null | Do not apply the update. |
| nonnegative_ accumulate | Add the update value to the current value, and set to 0 if the result is negative. |

Table S11: Dividers available in vivarium-core. Dividers are methods by which a variable's value is divided when division is triggered. New dividers can be easily defined and passed into a port schema (see Listing S4).

| Name | Function |
|---|---|
| set | The default divider. Daughters get the same value as the mother. |
| binomial | Sample the first daughter's value from a binomial distribution of the mother's value, and the second daughter gets the remainder. |
| split | Divide the mother's value in two. Odd integers will make one daughter receive 1 more than the other daughter. |
| split_dict | Splits a dictionary of $key : value$ pairs, with each daughter receiving a dictionary with the same keys, but with each value split. |
| zero | Daughter values are both set to 0. |
| no_divide | Asserts that this value should not be divided. |

#### 4.6 Code examples

This supplementary subsection includes some code listings demonstrating more advanced methods for specifying models with Vivarium.

Listing S4: Declaring custom updaters and dividers in `port_schema`. Updaters are functions that take a `current_value` and a `update_value`, and return the new value. Dividers are functions that take a `mother_value`, and a `state`, and return two values in a list – one for each daughter.

```
# updater that returns a random value
def random_updater(current_value, update_value):
    return random.random()

# divider that returns a random value for each daughter
def random_divider(mother_value, state):
    return [
        random.random(),
        random.random()]

def port_schema(self):
    ports = {
        'port1': {
            'variable1': {
                '_default': 1.0
                '_updater': {
                    'updater': random_updater
                    }
                '_divider': {
                    'divider': random_divider
            }
        }
    }
    return ports
```

Listing S5: Using glob ('*') schema to declare expected sub-store structure. In this example, `port1` is connected to sub-stores specified by a glob schema. This allows the process to read anything that port1 connects to which adheres to its declared schema. Sub-stores can be added and removed during runtime, and the process will see it.

```
def port_schema(self):
    ports = {
        'port1': {
            '*': {
                '_default': 1.0
            }
        }
    }
    return ports
```

Listing S6: Declaring process-store connections with `generate_topology`. A topology is a Python dictionary with keys for processes, and subkeys for their ports which map to paths at which they will connect to stores. A flat network does not require a path, just a store name at the same level. The syntax used for declaring paths is a Unix-style tuple, with every element in the tuple going further down the path from the root compartment, and '..' moving up a level to an outer compartment.

```
def generate_topology(self, config):
    topology = {
        'process': {
            'port1': ('path','to','store'), # connect port1 to inner compartment
            'port2': ('..','outer_store') # connect port2 to outer compartment
        }
    }
    return topology
```

Listing S7: Splitting a port into multiple stores with `generate_topology`. Variables read through the same port can come from different stores. To do this, the port is mapped to a dictionary with a `_path` key that specifies the path to the default store. Variables that need to be read from different stores each get their own path in that same dictionary. This same approach can be used to remap variable names, so different processes can use the same variable but see it with different names.

```
def generate_topology(self, config):
    topology = {
        # split a port into multiple stores
        'process1': {
            'port': {
                '_path': ('path_to','default_store'),
                'rewired_variable': ('path_to','alternate_store')
            }
        }
        # mapping variable names in process to different name in store
        'process2': {
            'port': {
                '_path': ('path_to','default_store'),
                'variable_name': 'new_variable_name'
            }
        }
    }
    return topology
```

Listing S8: Store API

```
# create the root
store = Store({})

# create a new store at a path
store.create(['top', 'store1'])
store.create(['top', 'store2'])

# create a process at a path
store.create(['top', 'process1'], ToyProcess({}))
```

```
store.create(['top', 'process2'], ToyProcess({}))

# connect port using a relative path
store['top', 'process1'].connect('port1', 'store1')

# connect using store target through a different port
store['top', 'process1'].connect('port2', store['top', 'process1', 'port1'])

# connect using absolute path
store['top', 'process1'].connect('port2', ('top', 'store2'), absolute=True)
```
